# The inducible response of the nematode *Caenorhabditis elegans* to members of its natural microbiome across development and adult life

**DOI:** 10.1101/554758

**Authors:** W Yang, C Petersen, B Pees, J Zimmermann, S Waschina, P Dirksen, P Rosenstiel, A Tholey, M Leippe, K Dierking, C Kaleta, H Schulenburg

## Abstract

The biology of all organisms is influenced by the associated community of microorganisms. In spite of its importance, it is usually not well understood how exactly this microbiome affects host functions and what are the underlying molecular processes. To rectify this knowledge gap, we took advantage of the nematode *C. elegans* as a tractable, experimental model system and assessed the inducible transcriptome response after colonization with members of its native microbiome. For this study, we focused on two isolates of the genus *Ochrobactrum*. These bacteria are known to be abundant in the nematode’s microbiome and are capable of colonizing and persisting in the nematode gut, even under stressful conditions. The transcriptome response was assessed across development and three time points of adult life, using general and *C. elegans*-specific enrichment analyses to identify affected functions. Our assessment revealed an influence of the microbiome members on the nematode’s dietary response, development, fertility, immunity, and energy metabolism. This response is mainly regulated by a GATA transcription factor, most likely ELT-2, as indicated by the enrichment of (i) the GATA motif in the promoter regions of inducible genes and (ii) of ELT-2 targets among the differentially expressed genes. We compared our transcriptome results with a corresponding previously characterized proteome data set, highlighting a significant overlap in the differentially expressed genes and the affected functions. Our analysis further identified a core set of 86 genes that consistently responded to the microbiome members across development and adult life, including several C-type lectin-like genes and genes known to be involved in energy metabolism or fertility. We additionally assessed the consequences of induced gene expression with the help of metabolic network model analysis, using a previously established metabolic network for *C. elegans*. This analysis complemented the enrichment analyses by revealing an influence of the *Ochrobactrum* isolates on *C. elegans* energy metabolism and furthermore metabolism of specific amino acids, fatty acids, and also folate biosynthesis. Our findings highlight the multifaceted impact of naturally colonizing microbiome isolates on *C. elegans* life history and thereby provide a framework for further analysis of microbiome-mediated host functions.

## Introduction

All multicellular organisms live in close association with microbial communities, the so-called microbiome or microbiota (McFall-Ngai et al., 2013). The microbiome appears to influence various biological functions of the host, for example food digestion and metabolism (Nicholson et al., 2012), development (Sampson and Mazmanian, 2015), immune defense (Tilg and Moschen, 2015), and aging (Heintz and Mair, 2014). It is also often linked to disease, such as obesity (Shen et al., 2013), liver cirrhosis (Abu-Shanab and Quigley, 2010), and even cancer (Garrett, 2015). However, the microbiome’s involvement in determining host traits is often based on correlation and exact information on its causal effects is usually absent due to missing possibilities for experimental manipulation. Well-established animal models may allow for such manipulations. Previous work with the fruitfly *Drosophila melanogaster* has indeed used an experimental approach to dissect the microbiome’s influence on host functions. These studies revealed that the microbiome influences gut morphology and physiological functions through changes in epithelial renewal rate, cellular spacing, and the distribution of different cell types in the epithelium (Broderick et al., 2014), that host genotype interacts with the microbiome to determine *Drosophila* nutritional state, as inferred from lipid content (Chaston et al., 2016), and that specific microbiome members enhance development and increase fly fitness (Pais et al., 2018). Moreover, microbiome members were demonstrated to interact with each other to influence host behavior, specifically olfactory and egg laying behaviors, mediated mainly through the odorant receptor *Or42b* (Fischer et al., 2017).

Another widely used model organism is the nematode *Caenorhabditis elegans*. The composition of its native microbiome has only been characterized relatively recently (Berg et al., 2016a; Dirksen et al., 2016; Samuel et al., 2016; Zhang et al., 2017). It includes a species-rich community of mainly Gammaproteobacteria and Bacteriodetes, including taxa from the genera *Enterobacter, Pseudomonas*, and *Ochrobactrum*. Most of the associated bacteria can be cultivated and are hence available for experimentation. Some of the cultivable isolates were already used to demonstrate their influence on *C. elegans* fitness under stress conditions, for example changes in nematode population growth under high osmolarity or temperature stress (Dirksen et al., 2016). Moreover, several distinct bacteria such as isolates of the genera *Pseudomonas, Enterobacter*, and *Gluconobacter*, were identified to enhance *C. elegans*’ immune defense against pathogens (Berg et al., 2016b; Dirksen et al., 2016; Montalvo-Katz et al., 2013; Samuel et al., 2016). To date, it is yet unclear how exactly the microbiome affects host molecular mechanisms to influence *C. elegans* life history characteristics.

The objectives of the current study were to fill this knowledge gap and obtain first insights into the effects of microbiome representatives on *C. elegans* molecular processes. For this study, we focused on two microbiome members of the genus *Ochrobactrum*, the isolates MYb71 and MYb237. These bacteria show the particular ability to colonize the nematode gut, even under adverse environmental conditions, and thereby form persistent associations with *C. elegans* (Dirksen et al., 2016). To assess the bacteria’s influence on the host, we performed a whole-genome transcriptome analysis across nematode development and adult life, using RNA sequencing. We used complementary types of enrichment analysis, including usage of a *C. elegans*-specific gene expression database (i.e., WormExp; (Yang et al., 2015b)) and an assessment of over-represented transcription factor binding sites (Shi et al., 2011), in order to characterize the affected biological functions and the signaling processes likely involved. We further compared our new data with a recently published proteome analysis of related material (Cassidy et al., 2018), in order to evaluate whether the inducible transcriptome and proteome vary, possibly indicating modifications after gene transcription. We further used the expression data for a reconstruction of the inducible metabolic activities in the worm gut, using the recently established metabolic network for *C. elegans* (Gebauer et al., 2016).

## Material and Methods

### *C. elegans* and bacteria strains

The canonical *C. elegans* laboratory strain N2 was used for all assays and generally maintained following standard procedures (Stiernagle, 2006). N2 was originally obtained from the CGC (Caenorhabditis Genetics Center), which is funded by NIH (National Institutes of Health) Office of Research Infrastructure Programs (P40 OD010440).

Three bacterial strains were used. The two Gram-negative bacteria *Ochrobactrum anthropic* strain MYb71 and *Ochrobactrum pituitosum* strain MYb237 are members of the native microbiome of *C. elegans*, obtained from the *C. elegans* isolate MY316 collected from a rotten apple in Kiel, Germany (Dirksen et al., 2016). The bacteria were freshly thawed for each experiment and cultured for two days at 25 °C on tryptic soy agar (TSA). Fresh colonies were used to produce liquid cultures in tryptic soy broth (TSB) at 28 °C in a shaking incubator for approximately 42 h. The *Escherichia coli* strain OP50 was used as a control and cultured in TSB at 37 °C in a shaking incubator overnight.

### Transcriptomic analysis of the inducible *C. elegans* response by RNA-Seq

N2 worms were maintained for a minimum of two generations on peptone-free medium (PFM) plates inoculated with the bacteria used for the experiments in order to allow adjustment to the bacteria. The transcriptome of *C. elegans* N2 was analyzed for six time points to cover various life stages: 6 h (second larval stage, L2), 24 h (L3), 48 h (L4), 72 h (1-day old adults, Ad1), 120 h (3-day old adults, Ad3) and 216 h (7-day old adults, Ad7). The worms were grown on 9 cm PFM plates with a 700 μl bacterial lawn (OD_600_ 10) of either MYb71, MYb237, or *E. coli* OP50. The worm stage was synchronized by bleaching. For each replicate 500 to 2000 synchronized hermaphrodites at the first larval stage (L1) were pipetted onto the bacterial lawn. The worms were maintained at 20 °C and harvested after the indicated periods. To separate the initially added worms from their offspring and to ensure sufficient food, worms were transferred to new plates every two days starting from first day of adulthood. Worm stages which did not produce eggs (L2, L3, and L4) were washed from the plates with M9 buffer and centrifuged to obtain a worm pellet. The supernatant was removed and 700 μl TRIzol™ (Thermo Fisher Scientific, Waltham, Massachusetts, USA) was added. Adult worms of Ad1, Ad3, and Ad7 were picked directly into 700 μl TRIzol™ to separate the initially placed worms from their offspring. All worms in TRIzol™ were five times frozen in liquid nitrogen and thawed at 46 °C in a thermo shaker to break open the cuticle. Subsequently, the samples were frozen and stored at −80 °C until the total RNA was extracted using the NucleoSpin RNA Kit (Macherey-Nagel, Düren, Germany). The transcriptome was analyzed for three replicates from independent runs of the exposure experiment. The only exception refers to L2 nematodes exposed to MYb237, for which only two independent replicates had sufficient amounts of RNA. All assays were performed without current knowledge of strain identity, and all treatment combinations were evaluated in parallel and in randomized order to avoid observer bias. RNA libraries were prepared for sequencing using standard Illumina protocols. Libraries were sequenced on an Illumina HiSeq™ 2000 sequencing machine with paired-end strategy at read length of 100 nucleotides. The raw data is available from the GEO database (Barrett et al., 2012; Edgar et al., 2001) under the GSE number GSE111364.

After removal of adaptor sequences and low quality reads via Trimmomatic (Bolger et al., 2014), RNA-Seq reads were mapped to the *C. elegans* genome (Wormbase version WS235; www.wormbase.org) by STAR 2.5.3a (Dobin et al., 2013) under default settings. Transcript abundance (read counts per gene) was extracted via HTSeq (Anders et al., 2015). Differential expression analysis was performed by aFold from ABSSeq (Yang et al., 2016). The log2 transformed fold-changes (*Ochrobactrum* vs. *E. coli* OP50) were taken as input for K-means cluster analysis using cluster 3.0 (de Hoon et al., 2004) with 8 initial clusters. A heat map was generated by TreeView version 1.1.4r3 (Saldanha, 2004). Core *Ochrobactrum* responsive genes were detected via aFold using a linear model to account for variation due to development and different adult life stages.

### Gene ontology, gene set, and motif enrichment analysis

Gene ontology (GO) analysis was performed using DAVID with a cut-off of FDR < 0.05 (Huang et al., 2009). A taxon-specific gene set enrichment analysis was performed using WormExp (Yang et al., 2015b), a web-based analysis tool for *C. elegans*, containing all of the available gene expression data sets for this nematode, thus allowing characterization of species-specific expression patterns. Only gene sets with FDR < 0.05 were considered to be significant. Motif analysis was carried out on the promoter regions, −600 bp and 250 bp relative to transcription start sites (TSS), of genes in each group. *De novo* motif discovery was performed using AMD (Shi et al., 2011).

### Transcriptome-proteome comparison

We compared our transcriptome results with the corresponding, previously published proteome data employing an isobaric labeling / LC-MS approach (Cassidy et al., 2018). The proteome data was generated from exposure experiments that were performed in almost identical form than those used for the transcriptome analysis. The main differences were that for the proteomics approach worms were grown on 15 cm plates (instead of 9 cm plates), Merck filters were used to separate larvae from the focal nematodes (instead of individual transfer of the focal worms with the help of worm-pickers), and only a single time-point was included, namely the young adult stage (i.e., after 72 h exposure of worms to the bacteria).

### Metabolic network analysis

Context-specific metabolic networks based on transcriptomic and proteomic data were reconstructed as described previously (Gebauer et al., 2016). Briefly, we used a two-step procedure in which first gene expression states were binarized into *on* and *off* and subsequently these states were used to derive activity of metabolic pathways through mapping to a genome-scale reconstruction of *C. elegans* metabolism using the iMAT procedure (Zur et al., 2010). In the first step, the procedure uses false discovery rate-adjusted p-values and fold-changes from differential expression analyses of the transcriptomic data to derive for each time point and condition the most likely activity state of a gene. While in the original procedure (Gebauer et al., 2016) this only involved comparisons to adjacent time points, we extended the approach by also comparing, for each time point, gene expression between all conditions. Differential gene expression analysis was performed using DESeq2 with standard parameters (Love et al., 2014). Only genes for which at least one comparison yielded an adjusted p-value below 0.05 across all comparisons were considered for the second step. The expression state of all other genes was left open in the second step of the analysis. In the second step, the iMAT-procedure (Zur et al., 2010) is used to derive a context-specific metabolic network that obeys the constraints of the network (steady state, flux bounds), maximizes the utilization of reactions associated with genes determined as *on* in the first step and minimized the utilization of reactions associated with genes determined as *off*. To determine changes in the activity of metabolic pathways, we used the subsystem annotation of each reaction present in the network reconstruction and counted the number of active reactions associated with each pathway for each condition and time point.

We next identified key enzymes involved in the response to bacterial colonization, using the metabolic transformation algorithm (MTA, (Yizhak et al., 2013)). The metabolic transformation algorithm is able to identify reaction knockouts that are best able to transform a metabolic network from a given source state to a desired target state. We modified MTA in two points. First, we used the context-specific metabolic networks we reconstructed as described above as source state instead of the purely iMAT-derived metabolic networks used in the original approach (Yizhak et al., 2013) to maximize comparability of the identified key enzymes to the context-specific metabolic networks we derived. Second, to facilitate interpretation, we did not consider single-reaction knockouts but rather gene knockouts. Thus, we did not assess the ability of knockouts of specific reactions to move the source network towards the target state but rather we performed knockouts on the level of genes. Thus, we tested for each metabolic gene present in the reconstruction whether its knockout blocked any reaction and performed MTA on the metabolic network after constraining the flux through all correspondingly blocked reactions to zero.

We performed two sets of MTA runs on the transcriptomic and the proteomic data: one for the transition between growth of either *Ochrobactrum* species to growth on *E. coli* OP50 and vice versa. For each set of runs we performed MTA runs for every time point and each comparison. Thus, for the analysis of the metabolic transition between growth on *Ochrobactrum* to growth on *E. coli* OP50, we considered for each time point either of the two *Ochrobactrum*-growth specific metabolic networks as source state and the *E. coli* OP50-specific metabolic network as target state. MTA returns for each gene a score indicating to which extent the knockout of this gene shifts the source state towards the target state. Moreover, MTA determines a threshold score at which the knockout of a gene is considered to lead to a significant shift towards the target state. Since the absolute values of MTA-scores change between runs (but rarely their order), we summarized each run by setting all runs with an MTA-score equal or below the cut-off threshold to zero, replacing the remaining scores with their rank in the ordered list and normalizing values to a maximum of 1 (highest MTA score) and a minimum of 0. For each comparison we performed five bootstrap runs in which we randomly removed 10 % of gene expression values before performing MTA to assess robustness of results. MTA-scores were summarized through averaging the rank-normalized scores for each gene across all runs of a set and subsequent normalization of scores to a range from 0 to 1. Subsequently, an overall MTA score for each gene was determined by summing MTA-scores for the transition from growth on *Ochrobactrum* to growth on *E. coli* OP50 using transcriptomic and proteomic data and subtracting the scores for the opposite transition. As a result, we obtained an overall MTA score which is positive if a gene mediates the transition from growth on *Ochrobactrum* to growth on *E. coli* OP50 and negative in the other direction.

## Results and Discussion

### Transcriptional variation is determined by nematode development and microbial exposure

The *C. elegans* microbiome isolates *Ochrobactrum* MYb71 and *Ochrobactrum* MYb237 are able to efficiently colonize the nematode gut (Figure 1A and also (Dirksen et al., 2016)). To assess the effects of MYb71 and MYb237 on molecular processes in the *C. elegans* host, we performed transcriptome analysis using RNAseq across three developmental stages (L2, L3, and L4) and three time points during adulthood (day 1, day 3, and day 7). The standard laboratory food bacterium *Escherichia coli* OP50 served as control (Figure 1B). We first used principal component analysis (PCA) to explore transcriptomic variation across treatments and time points. We found that both the first and second principal component separate different *C. elegans* developmental stages (Figure 1C), suggesting development as the main determinant of gene expression variation. The different bacterial treatments account for less overall variation. As indicated by the third principal component, it appears that colonization by the *Ochrobactrum* strains and the *E. coli* control produce more pronounced differences at the middle time points (i.e., L4 larval stage and the first adult time point Ad1; Figure 1D). This observation may suggest that *Ochrobactrum* and *E. coli* vary in how they affect nematode development from L4 to adult.

**Figure 1.**
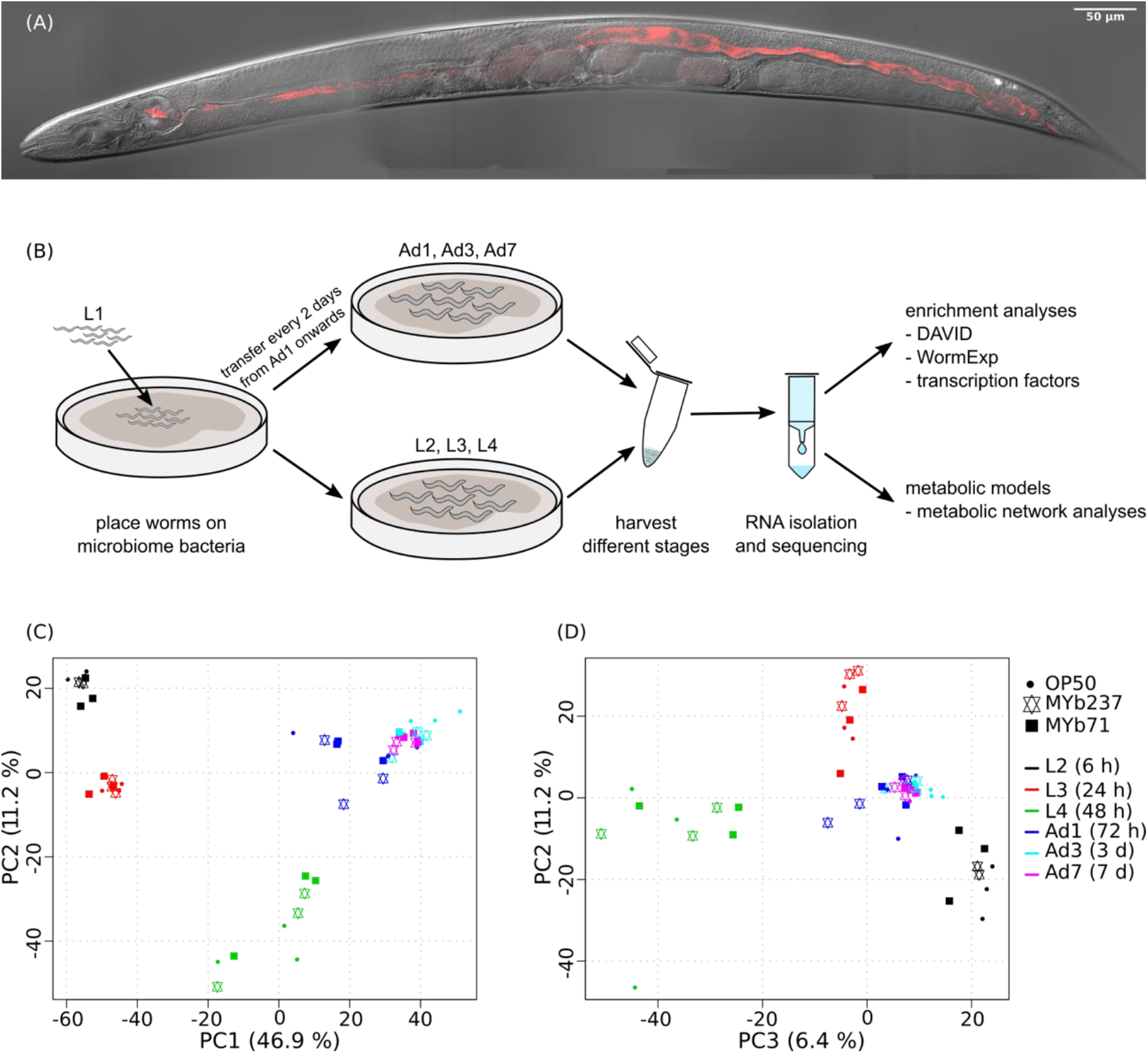
Colonized *C. elegans* individual, workflow and principal component analysis of transcriptomic variation. (A) Colonization of *C. elegans* strain N2 by Ochrobactrum MYb71, highlighting distribution of the microbiome isolate (labeled with red fluorescence) throughout the pharynx and gut lumen. (B) Workflow. (C, D) Variation was assessed for N2 fed with *E. coli* OP50 (indicated by filled cycle) or colonized by *Ochrobactrum* isolates MYb237 (triangles up and down) or MYb71 (filled squares) at six time points including the second larval stage (L2, 6 h), L3 (24 h), L4 (48 h), 1-day old adults (Ad1), Ad3 (3 d), and Ad7 (7 d), as indicated by different colors.

Next, we specifically assessed transcriptional variation by either *Ochrobactrum* strains *versus E. coli* for each time point, in order to explore the gene functions affected by these microbiome members. A total of 893 genes were differentially expressed, falling into eight clusters of co-regulated genes, as identified through K-means clustering (Figure 2A). These observations highlight that the transcriptional response is indeed influenced by *Ochrobactrum* across time, whereby the two *Ochrobactrum* strains do not appear to vary much in inducible expression patterns (Figure 2A). Clusters 1 and 5 refer to genes with strong differential expression across all time points (up- and down-regulated genes, respectively), while other clusters show differential expression at specific time points only. This pattern possibly indicates that *Ochrobactrum* influences *C. elegans*’ life history in more than one way.

**Figure 2.**
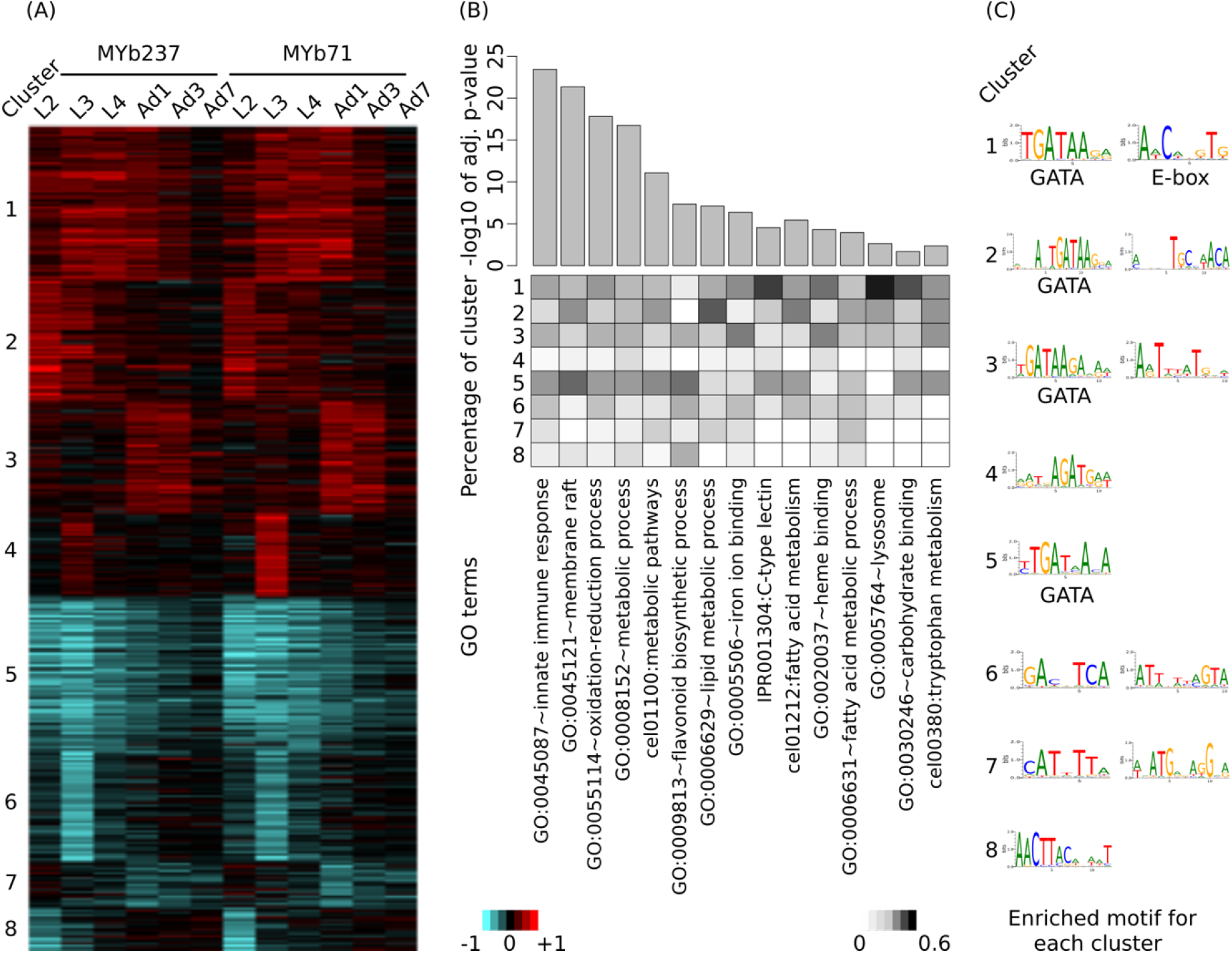
Induced differential gene expression in *C. elegans* after colonization with *Ochrobactrum* isolates. Differential expression is identified via comparison of worms colonized by the *Ochrobactrum* isolates *versus* those fed with *E. coli* OP50. (A) Co-regulation of differentially expressed genes. Eight clusters (indicated by the numbers on the left) of co-regulated genes were identified via K-means clustering. Red and blue colors refer to up and down regulation, respectively. Heatmap scale bars indicate fold changes at log_2_ scale. (B) Enriched gene ontology (GO) terms of differentially expressed genes. GO enrichment analysis was performed by DAVID. 15 selected GO terms are shown with significance (top panel, bar plot with adjusted p-values). The percentage of genes, which contributed from each cluster to the identified GO terms is shown as a heatmap in the middle panel. (C) Enriched transcription factor binding sites (Motif) for each cluster. Motifs were detected by AMD at the promoter region of genes.

To uncover the affected biological processes, we applied gene ontology (GO) enrichment analysis on all identified 893 differentially expressed genes. A variety of GO terms were found to be significantly over-represented among these genes (Supplementary Table S1). Several of the most significant GO terms are related to immunity, for example GO:0045087 (e.g., innate immune response) and IPR001304 for the C-type lectins (Pees et al., 2015), the latter term particularly over-represented among the up-regulated gene clusters (Figure 2B). These results are consistent with repeated previous reports of a link between the microbiome and the host immune system (Cullender et al., 2013; Thaiss et al., 2016; Tilg and Moschen, 2015). Moreover, many metabolism-related processes are significantly enriched (e.g., GO:0055114∼oxidation-reduction process and GO:0006629∼lipid metabolic process; Figure 2B; Supplementary Table S1). This may suggest an influence of *Ochrobactrum* on *C. elegans* metabolism, consistent with previous work on the alternative food bacterium *Comamonas* DA1877 (MacNeil et al., 2013). Interestingly, clusters 1 and 5 show a stronger relationship to the indicated GO terms than clusters related to only one time point. This may imply that the major effects of these microbiome members on *C. elegans* persist across the various life stages.

The complementary enrichment analysis with WormExp (Yang et al., 2015b) revealed significant over-representation of gene sets, which were related to pathogen infection (e.g., nematocidal *Bacillus thuringiensis* strain BT247 (Yang et al., 2015a), *Staphylococcus aureus* (Bond et al., 2014), *Pseudomonas aeruginosa* strain UCBPP-PA14 (Nakad et al., 2016); Table 1, Supplementary Table S1). Moreover, data sets related to the GATA transcription factor gene *elt-2* (RNAi (Schieber and Chandel, 2014), ChIP-Seq targets (Mann et al., 2016)) or the E-box transcription factor gene *hlh-30* (study on *S. aureus* (Visvikis et al., 2014) and *hlh-30* mutant (Grove et al., 2009)) were also enriched, suggesting an involvement of these transcription factors in the nematode’s response to the microbiome. Additional over-represented gene sets relate to *C. elegans* dietary responses, for example gene sets related to the insulin-like pathway (*daf-2* (Knutson et al., 2016) and *daf-16* (McElwee et al., 2004)), fasting (Lee et al., 2014, 1), and starvation (Mueller et al., 2014). We further found an enrichment of the worm’s response to the previously studied food bacterium *Comamonas* DA1877 (MacNeil et al., 2013). Overall, our results suggest that the *Ochrobactrum* isolates affect dietary responses, metabolic processes, and also interact with the immune system.

**Table 1.** Enriched WormExp gene sets for differentially expressed genes upon *Ochrobactrum* colonization.

| Category | Gene set | Counts | Bonferroni |
| --- | --- | --- | --- |
| Microbes | down by *B. thuringiensis* at 6 h (BT247, 1:10) (Yang) | 232 | 4.13E-149 |
| TF Targets | low-complexity *elt-2* targets | 338 | 4.68E-133 |
| DAF/Insulin/food | down by *daf-2* mutant (Knutson) | 171 | 7.24E-130 |
| Mutants | down by *elt-2* RNAi under normoxia | 141 | 4.78E-90 |
| Microbes | up by PA14, 12 h | 106 | 1.29E-81 |
| DAF/Insulin/food | down by fasting (Lee) | 90 | 8.03E-65 |
| Microbes | down by Bt toxin, Cry5B | 110 | 7.36E-64 |
| Microbes | down by PA14, 12 h | 63 | 7.59E-62 |
| Microbes | down by *S. aureus* (Bond) | 84 | 2.05E-53 |
| Microbes | down by *B. thuringiensis* (BT247), 12 h | 157 | 2.32E-49 |
| DAF/Insulin/food | down by starved (Mueller) | 170 | 4.55E-48 |
| Microbes | down fed by *E. coli* HT115 vs. OP50 | 38 | 3.25E-47 |
| DAF/Insulin/food | up by *daf-16* mutant (McElwee) | 90 | 1.84E-46 |
| Microbes | up by *B. thuringiensis* at 6 h (BT247, 1:10) (Yang) | 112 | 2.35E-41 |
| DAF/Insulin/food | down by *daf-16* mutant (McElwee) | 95 | 3.11E-39 |
| Microbes | up by *S. aureus*, dependent on *hlh-30* (Visvikis) | 100 | 1.07E-38 |
| Microbes | up by *S. aureus* (Bond) | 85 | 5.18E-36 |
| DAF/Insulin/food | down fed by *Comamonas* DA1877 vs. OP50 young adult | 67 | 8.49E-33 |
| DAF/Insulin/food | up by fasting (Lee) | 52 | 1.39E-25 |
| DAF/Insulin/food | up by starved (Mueller) | 140 | 3.55E-25 |
| DAF/Insulin/food | up fed by *Comamonas* DA1877 vs. OP50 young adult | 31 | 1.06E-23 |
| Mutants | up by *elt-2* RNAi under normoxia | 39 | 3.64E-18 |
| Microbes | up fed by *E. coli* HT115 vs. OP50 | 19 | 1.21E-16 |
| DAF/Insulin/food | up by *daf-2* mutant (Knutson) | 41 | 2.90E-13 |
| Microbes | up by Bt toxin, Cry5B | 46 | 4.86E-12 |
| Mutants | down *hlh-30* mutant | 27 | 7.65E-12 |

The presence of co-regulated gene clusters, as inferred through our cluster analysis, could be caused by transcription factors. To explore this idea, we performed *de novo* motif enrichment analysis on promoter regions of the genes within each cluster. We identified one or two informative transcription factor binding motifs for each cluster (Figure 2C). However, only two of them have been previously characterized for *C. elegans*: the GATA motif with consensus sequence GATAA and the E-box with the sequence motif CACGTG. GATA transcription factors are known to play a role in immunity (Shapira et al., 2006), intestine development (mainly through ELT-2 (McGhee et al., 2007)), and aging (mainly through ELT-3, ELT-5, and ELT-6 (Budovskaya et al., 2008)). The GATA motif is enriched in clusters 1, 2, 3, and 5, which generally showed an over-representation of GO terms and gene sets related to above characteristics, especially immunity (Figure 2B, Table 1). E-box transcription factors were shown to shape *C. elegans* immunity (e.g., *hlh-30* (Visvikis et al., 2014)) and muscle development (Grove et al., 2009). The corresponding motif was only identified for cluster 1, which similarly produced an enrichment for immunity-related GO terms. As the motif analysis is corroborated by the enriched gene expression sets for mutants of known GATA and E-box transcription factors, we conclude that these regulators play a central role in coordinating the response of *C. elegans* to its microbiome members.

### Signature genes in the *C. elegans* response to *Ochrobactrum*

To identify genes specific for the response to the microbiome members only, we employed a model-based statistical analysis (based on the linear model-option in aFold (Yang et al., 2016)), in which we statistically accounted for the factor time. Thus, these genes specifically respond to the presence of *Ochrobactrum*, irrespective of any variation in gene expression across worm development. This analysis revealed a total of 86 differentially regulated genes: 65 differentially expressed in the presence of MYb237 and 71 in the presence of MYb71. More than 70 % of these genes were identical, confirming our above notion that these two *Ochrobactrum* isolates induce a similar expression response. Of the total of 86 genes, 18 and 59 were continuously up- and down-regulated across the six studied time points, respectively (Figure 3A). Of these genes, six showed a more than twofold change in expression. They may therefore be suited as indicator markers for the *C. elegans* response towards *Ochrobactrum* colonization. They include the four down-regulated genes *acdh-1, metr-1, cth-1*, and *mtl-2*, and the two up-regulated genes Y53G8AM.5 and F59D6.3. *acdh-1* and *metr-1* were previously identified as reporters for a dietary response (MacNeil et al., 2013; Watson et al., 2013). The down-regulated gene *mtl-2* is known to play a role in regulating growth and fertility (Freedman et al., 1993). *cth-1* encodes a putative cystathionine gamma-lyase, which may contribute to metabolic processes.

**Figure 3.**
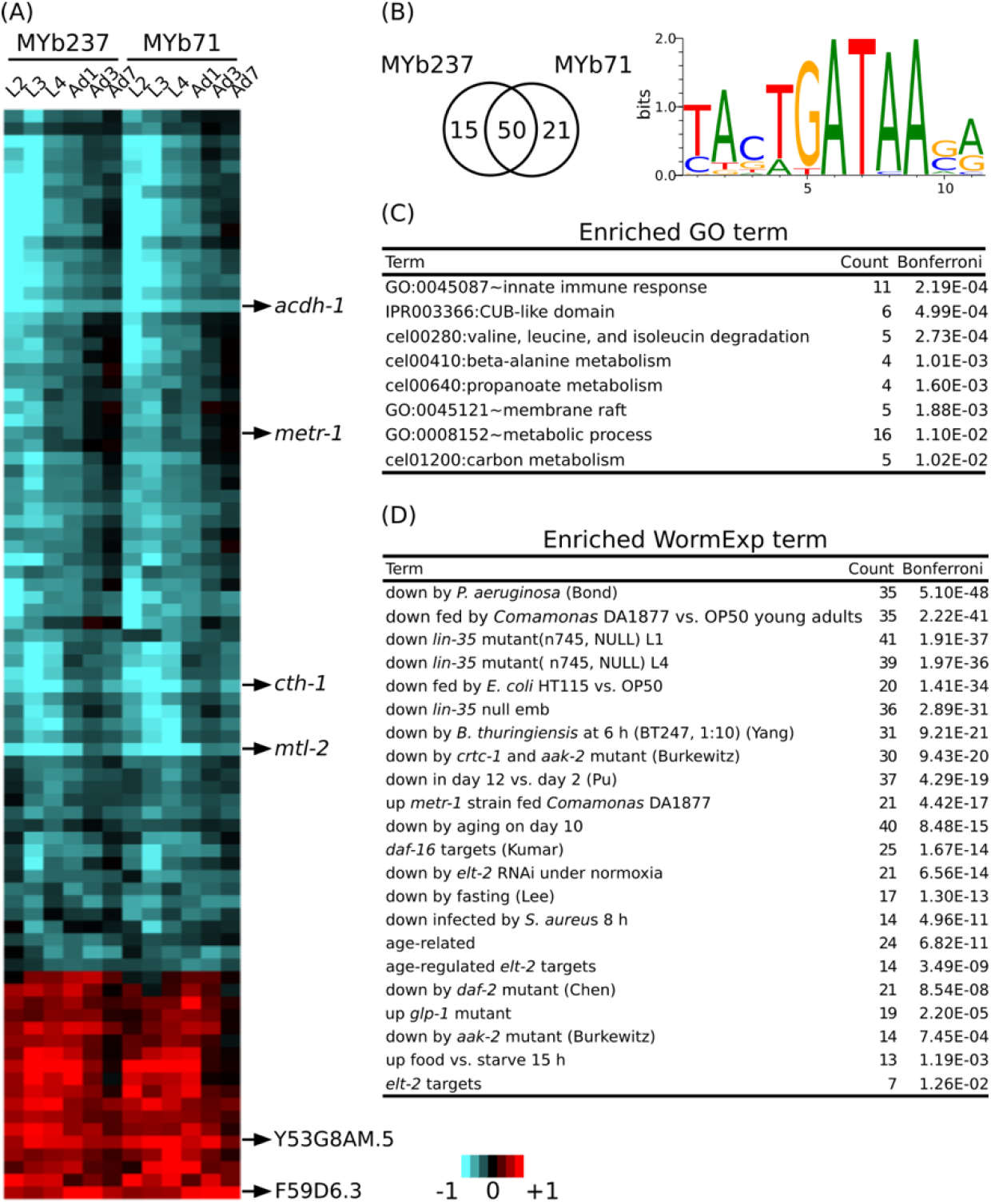
Core responsive genes. (A) Differential gene expression of 86 core responsive genes are shown using a heatmap. Genes with the strongest change of expression are marked on the right. Red and blue colors refer to up and down regulation, respectively. Heatmap scale bars indicate fold changes at log_2_ scale. (B) Venn diagram showing the overlap of core responsive gens induced upon colonization by MYb71 and MYb237 and the enriched binding motif. (C) Enriched GO terms. GO enrichment analysis was performed by DAVID. (D) Enriched WormExp gene sets.

A motif enrichment analysis on these 86 genes again suggests that the microbiome response is controlled by a GATA transcription factor (Figure 3B). This is additionally supported by the significantly over-represented gene set that is downstream of the GATA transcription factor gene *elt-2* (Figure 3C; Supplementary Table S2). Our enrichment analysis generally yielded results that are consistent with above analysis (cf. Figure 2B). They indicate a potential interaction between *Ochrobactrum* and immunity and also the involvement of the dietary response, the microbe’s influence on metabolic pathways, energy production, and the response to the bacterial food source *Comamonas* DA1877 (Figure 3C; Supplementary Table S2). Moreover, in this analysis, we also found enriched gene sets related to development (i.e., downstream targets of *lin-35* (Kirienko and Fay, 2007)) and fertility (i.e., downstream targets of *glp-1* (Gracida and Eckmann, 2013)). The latter suggest that the microbiome additionally affects these two characteristics in *C. elegans*.

### Concordance between the *C. elegans* proteome and transcriptome responses to *Ochrobactrum*

We next compared the transcriptome response at the first adult time point to a corresponding proteome data set, which was obtained for the same time point upon colonization with *Ochrobactrum* (Cassidy et al., 2018). This proteome data set revealed significant differential abundance of 123 out of more than 3,600 quantified proteins. Of these, 50 had higher and 73 had lower abundance. These differentially abundant proteins showed consistent changes at the transcript level upon colonization of nematodes to both MYb71 and MYb237 (Figure 4A, 4B). In detail, 40 genes showed consistently higher (80 % of the 50 up-regulated proteins) and 67 consistently lower abundance (91.8 % of the 73 down-regulated proteins) in both the transcriptome and proteome data sets. One example is the short-chain acyl-CoA dehydrogenase gene, *acdh-1*, which was consistently and strongly down-regulated in the presence of the two *Ochrobactrum* isolates. Similarly, four C-type lectin-like genes produced consistent and significant expression changes at transcript and protein levels (up-regulation: *clec-63* and *clec-65*, down-regulation: *clec-47* and *clec-218*). These results were in line with the enriched GO term of C-type lectin genes (Figure 2B), suggesting an important role of C-type lectins in mediating the nematode’s interaction with specific microbiome members, which are highly numerous in nematodes and could contribute to recognition of microbe-associated molecular patterns (MAMPs) (Pees et al., 2015). Only two genes showed significant opposite expression patterns at transcript and protein levels. These included two members of the *C. elegans* lysozyme gene family (Boehnisch et al., 2011), *lys-4* and *lys-5*, which were up-regulated at mRNA, yet down-regulated at protein level, indicating a post-transcriptional regulation.

**Figure 4.**
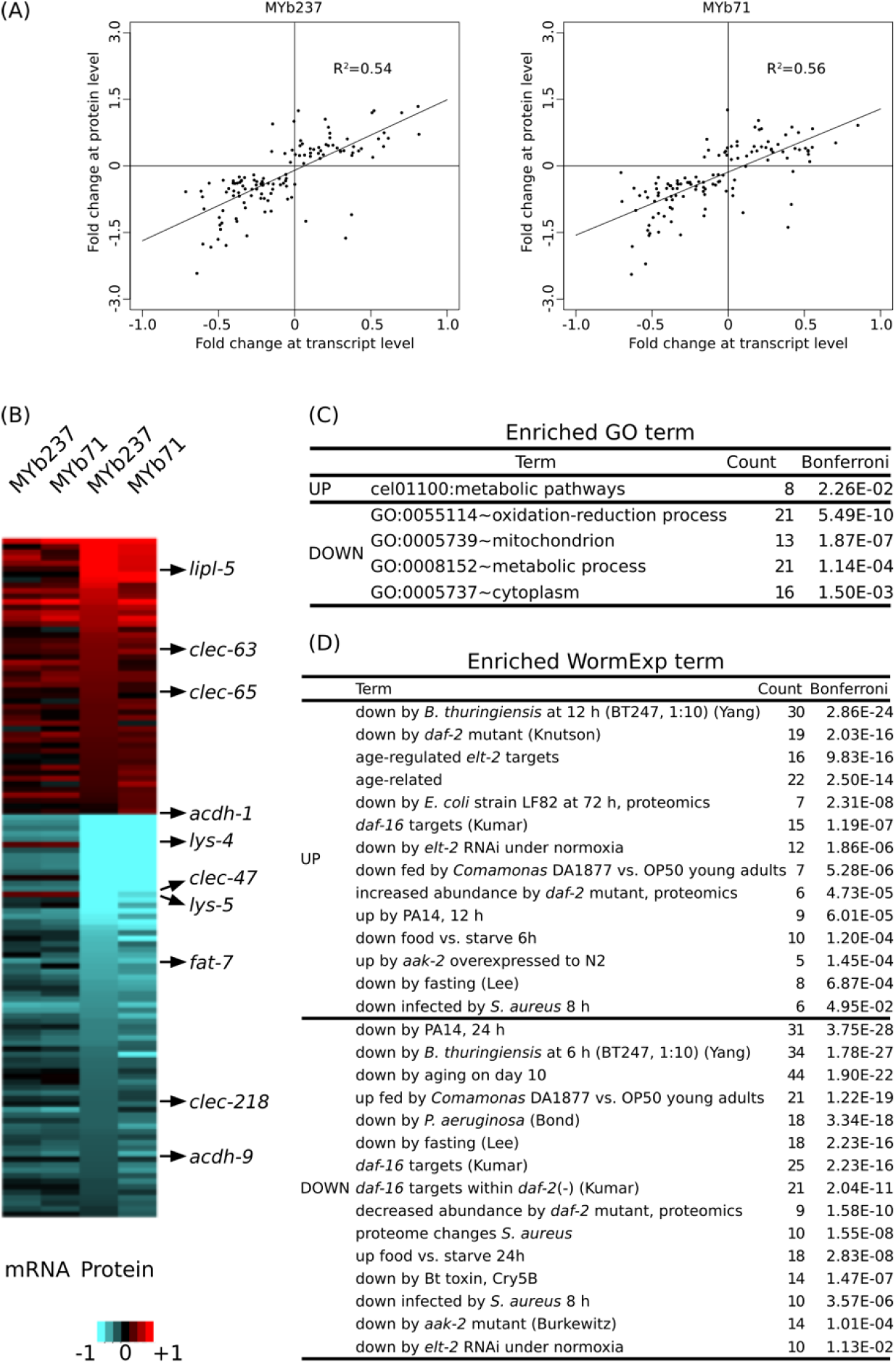
Comparison of differential gene expression at transcript and protein levels. (A) Comparison of expression levels at transcript and protein level. Data are only shown for genes with significant differential expression at protein level. Results are shown separately upon exposure to either MYb71 or MYb237. (B) Expression changes of genes at transcript and protein levels at the 1-day old adult stage (Ad1, 72 h). Red and blue colors refer to up and down regulation, respectively, or higher and lower protein abundances. Heatmap scale bars indicate fold changes at log_2_ scale. A selection of genes with functional information is shown on the right. (C) Enriched GO terms. GO enrichment analysis was performed by DAVID. (D) Enriched WormExp gene sets.

An enrichment analysis of the 123 differentially expressed proteins revealed several significantly enriched functions. These included GO terms related to metabolism and energy production, such as the terms cel01100:Metabolic pathways, GO:0008152∼metabolic process, and GO:0005739∼mitochondrion (Figure 4C; Supplementary Table S3). The enriched term GO:0055114∼oxidation-reduction process could similarly indicate a role in energy metabolism or, alternatively, stress response. The WormExp analysis yielded similar results as above, including significant over-representation of gene sets related to immunity, dietary response, ageing, and that controlled by *elt-2*. We also found enrichment of genes controlled by *aak-2*, which is part of the AMPK pathway (Burkewitz et al., 2015; Hou et al., 2016) and thus further supports the possible influence of the microbiome members on *C. elegans* energy production (Figure 4D).

### Metabolic network analysis indicates colonization-specific changes in fatty acid, amino acid, folate, and energy metabolism

Since we observed considerable metabolism-associated changes in response to bacterial colonization, we performed a more detailed analysis of metabolic changes associated to bacterial colonization. To this end, we derived context-specific metabolic networks by mapping the transcriptomic and proteomic data to a genome-scale reconstruction of *C. elegans* metabolism (Gebauer et al., 2016). This method comprises two steps that initially discretizes gene expression states into on and off based on differential expression between conditions. In the second step, a subnetwork of the metabolic network is determined that can carry flux and maximizes the utilization of reactions catalyzed by enzymes that are active based on the first step while minimizing the utilization of reactions catalyzed by enzymes that are inactive (Gebauer et al., 2016).

In a first step, we compared the reconstructed context-specific networks of the different developmental stages upon exposure to the different bacteria separately (Figure 5A). In agreement with the expression-based analysis, we found that developmental stage had the strongest impact on metabolic activity. Bacterial colonization affected metabolism mostly during larval development and had only little impact in adult worms. On a global scale, amino acid metabolism, carbohydrate metabolism, and vitamin metabolism were most strongly affected by bacterial colonization (Figure 5B, Supplementary Tables S4 and S5). Differences among bacterial effects were found for fatty acid metabolism, the metabolism of various amino acids (branched-chain amino acids, cysteine/methionine metabolism, tryptophan metabolism, lysine metabolism), and also folate (Figure 5C). Intriguingly, folate metabolism has previously been observed to be a key process involved in the modulation of host physiology and lifespan by food microbes (Cabreiro et al., 2013). Moreover, branched-chain amino acids, cysteine as well as methionine and tryptophan are important modulators of *C. elegans* nutritional and stress responses (Gebauer et al., 2016; Lee et al., 2015; Mansfeld et al., 2015; van der Goot et al., 2012).

**Figure 5.**
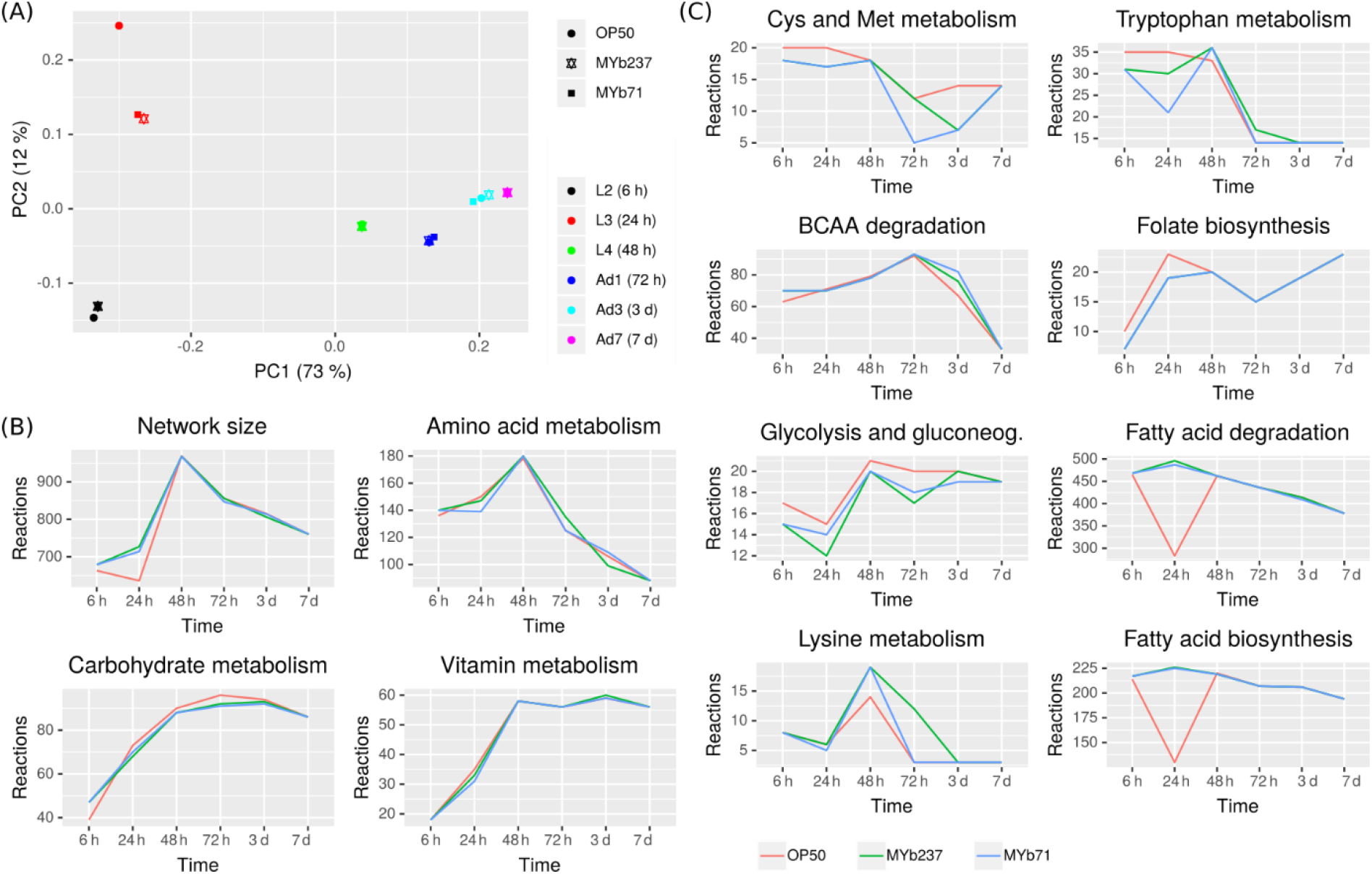
Metabolic response of *C. elegans* to bacterial colonization and exposure. (A) Principal component analysis of context-specific metabolic networks. (B) Changes in the number of active reactions in top-level pathways across developmental stages and bacterial treatment. (C) Changes in the number of active reactions in individual metabolic pathways. Only eight pathways with highest absolute difference in the number of active reactions between *Ochrobactrum* (MYb237 and MYb71) and *E. coli* OP50 exposure across all time points are shown. Numbers of active reactions in all subsystems are provided in Supplementary Tables S4 and S5.

In a second step, we used the metabolic transformation algorithm (Yizhak et al., 2013) to identify enzymes that most strongly contributed to the metabolic shifts associated with bacterial colonization (Supplementary Table S6). The metabolic transformation algorithm identifies knockouts in a metabolic network that shift the metabolic network from a specific source state to a desired target state. Thus, this algorithm allows us to assess which enzymatic processes mediate the metabolic transition from *C. elegans* growth on the *Ochrobactrum* strains towards growth on *E. coli* OP50 and *vice versa*. In order to ensure robustness of the results, we used combined data sets of the transcriptomic and proteomic results for these analyses. For the transition from *E. coli* to *Ochrobactrum* (i.e., requirements for growth on the microbiome bacteria), we observed a considerable number of potential key mediators in N-glycan biosynthesis, serotonin/octopamine biosynthesis and co-factor metabolism. For N-glycan biosynthesis, several genes in N-glycan precursor biosynthesis including *algn-1* – *algn-3, algn-5, algn-7* and *algn-10* – *algn-13* were identified. N-glycans have been implicated in the interaction with microbial pathogens in *C. elegans* (Butschi et al., 2010; Shi et al., 2006) and commensals in higher organisms (Koropatkin et al., 2012). Thus, our results may indicate a role of N-glycans also in the interaction of *C. elegans* with its microbiome members. For serotonin/octopamine metabolism, we identified tyrosine decarboxylase (*tdc-1*) as well as tyramine beta-hydroxylase (*tbh-1*) involved in octopamine biosynthesis and tryptophan hydroxylase (*tph-1*) involved in serotonin biosynthesis as key mediators of metabolic transitions. Serotonin and octopamine are antagonists that are important modulators of *C. elegans* behavior and energy metabolism (Niacaris and Avery, 2003; Noble et al., 2013; Suo et al., 2009). For the transition from *Ochrobactrum* to *E. coli* OP50 (i.e., requirement for growth on *E. coli*), we identified only two enzymes as key mediators: a predicted adenosine kinase, R07H5.8, and C05C10.3 involved in ketone body metabolism as well as the first step in mevalonate/terpenoid metabolism. Interestingly, adenosine kinase catalyzes the conversion of adenosine and ATP to AMP and ADP and thereby influences central aspects of *C. elegans* physiology via AMPK-signaling.

### Conclusions

We here provide the first study on the transcriptional response of *C. elegans* towards colonizing members of its native microbiome, both from the genus *Ochrobactrum*. This response was characterized across three larval and three adult stages, capturing different parts of the worm’s life cycle. We link the transcriptomic results with proteome data and further combine gene expression enrichment analyses with metabolic network modelling, thereby providing a comprehensive reference framework for the inducible molecular response of *C. elegans* towards colonizing microbiome members.

We found that there is a specific response to *Ochrobactrum* that involves several biological processes, such as those related to immunity, ageing, fertility, and development, and the three connected characteristics dietary response, metabolism, and energy production (Figure 2). We further identified an *Ochrobactrum*-exclusive signature of 86 differentially expressed genes (Figure 3), some of which are directly related to above functions and could serve as indicator genes for the worm’s response to this microbiome taxon. We additionally identified a significant overlap between the *Ochrobactrum*-mediated response at transcript and protein levels (Figure 4), thereby providing one of the few examples in *C. elegans* which could directly connect these two levels within one study set-up. This overlapping set of genes is again enriched in functions related to pathogen responses, dietary responses, and energy metabolism, highlighting that these are indeed central processes affected by the presence of colonizing microbiome members rather than non-colonizing food bacteria. Our metabolic network analysis confirms the influence of *Ochrobactrum* on energy metabolism (e.g., via the AMPK pathway; (Burkewitz et al., 2015; Hou et al., 2016)), and furthermore metabolism of specific amino acids, fatty acids, and also folate biosynthesis (Figure 5).

Our complementary analyses consistently identify a role of GATA transcription factors, most likely ELT-2, in coordinating the response to *Ochrobactrum* (Table 1; Figures 2C, 3B, 3D, 4D). ELT-2 is known to control gene expression in the intestine (McGhee et al., 2007) where the microbiome resides, thus providing ample opportunity for direct interactions between bacterial molecules and upstream regulators of ELT-2 and, conversely, between ELT-2 downstream factors and the colonizing bacteria. Moreover, ELT-2 was suggested to act as central regulator of inducible gene expression against pathogenic bacteria in the adult intestine (Block and Shapira, 2015; Yang et al., 2016b), thereby providing a link to the enriched set of immune-related genes. However, *Ochrobactrum* does not behave as a pathogen, as it neither enhances nematode mortality (compared to the standard food source *E. coli*) nor causes any obvious damage during intestinal colonization (Figure 1A). Therefore, the enrichment of immune-related genes may indicate multiple functions of the pathogen-responsive genes, which could be involved in both digestion and pathogen degradation or the recognition of any gut-colonizing microbe (e.g., possible for the enriched group of C-type lectins; (Pees et al., 2015)). This enriched category may also reflect the previous notion that immune-related genes are generally involved in coordinating composition of the host-associated microbes (Eberl, 2010; McFall-Ngai, 2007).

## Acknowledgements

We are grateful for comments and suggestions from members of the Schulenburg and Kaleta groups and the Collaborative Research Center CRC 1182 on Origin and Function of Metaorganisms, funded by the Deutsche Forschungsgemeinschaft (DFG; German Science Foundation). We acknowledge financial support by the CRC 1182 to HS (project A1.1), KD (project A1.2), ML (project A1.3), AT (project A1.4), CK (project INF), and PR (project Z3). CK, PR, and HS were additionally supported by the DFG under Germany’s Excellence Strategy – EXC 2167-390884018, and HS further by a fellowship from the Max-Planck Society.

## Author Contributions Statement

The study was designed by WY, CP, BP, AT, ML, KD, CK, and HS. CP, BP, and PD performed nematode experiments. CP isolated RNA. PR coordinated RNA sequencing. YW performed transcriptome and proteome analysis. JZ, SW, and CK reconstructed and analyzed metabolic network models. All authors discussed and interpreted the data. WY, BP, CP, CK, and HS wrote the manuscript.

## Supplementary Material

**Supplementary Table S1. Fold-change in expression of differentially expressed genes, gene ontology (GO) term enrichment, and WormExp enrichment.**

**Supplementary Table S2. Core responsive genes induced upon *Ochrobactrum* colonization, GO term enrichment, and WormExp enrichment.**

**Supplementary Table S3. Comparison of differential expression at transcript and protein levels and WormExp enrichment.**

**Supplementary Table S4: Inferred active reactions in metabolic subsystems across developmental stages and different bacterial microbiome species.**

**Supplementary Table S5: Inferred active reactions in individual metabolic pathways across developmental stages and different bacterial microbiome species.**

**Supplementary Table S6: Identification of key enzymes mediating the response to *Ochrobactrum* species and *Escherichia coli.***

**Supplementary Movie 1. 3D reconstruction of *Ochrobactrum* MYb71 colonization in a *C. elegans* N2 host via fluorescence *in situ* hybridization. Signal of general bacterial probe EUB338 is shown in red, nuclei stained with DAPI in blue. The individual shown is the same as in figure 1A.**

